# Sex-related differences in behavioral and amygdalar responses to compound facial threat cues

**DOI:** 10.1101/179051

**Authors:** Hee Yeon Im, Reginald B. Adams, Cody A. Cushing, Jasmine Boshyan, Noreen Ward, Kestutis Kveraga

## Abstract

During face perception, we integrate facial expression and eye gaze to take advantage of their shared signals. For example, fear with averted gaze provides a congruent avoidance cue, signaling both threat presence and its location, whereas fear with direct gaze sends an incongruent cue, leaving threat location ambiguous. It has been proposed that the processing of different combinations of threat cues is mediated by dual processing routes: reflexive processing via magnocellular (M) pathway and reflective processing via parvocellular (P) pathway. Because growing evidence has identified a variety of sex differences in emotional perception, here we also investigated how M and P processing of fear and eye gaze might be modulated by observer’s sex, focusing on the amygdala, a structure important to threat perception and affective appraisal. We adjusted luminance and color of face stimuli to selectively engage M or P processing and asked observers to identify emotion of the face. Female observers showed more accurate behavioral responses to faces with averted gaze and greater left amygdala reactivity both to fearful and neutral faces. Conversely, males showed greater right amygdala activation only for M-biased averted-gaze fear faces. In addition to functional reactivity differences, females had greater bilateral amygdala volumes, which positively correlated with behavioral accuracy for M-biased fear. Conversely, in males only the right amygdala volume was positively correlated with accuracy for M-biased fear faces. Our findings suggest that M and P processing of facial threat cues is modulated by functional and structural differences in the amygdalae associated with observer’s sex.

## Introduction

Face perception, particularly assessment of facial emotion during social interactions, is critical for adaptive social behavior. It enables both observers and expressers of facial cues to communicate nonverbally about the social environment. For example, a happy facial expression implies to an observer that either the expresser or the environment surrounding the observer is safe and friendly, and thus approachable, whereas a fearful facial expression can imply the existence of a potential threat to an expresser or even to an observer. Prior work has investigated how observers read such signals from a facial expression when it is combined with direct or averted eye gaze, which impart different meanings to a facial expression. It has been shown that a fearful face tends to be perceived as more fearful when presented with an averted eye gaze because the combination of fearful facial expression and averted gaze provides a congruent social signal (both the expression and the gaze direction signal avoidance), leading to facilitated processing of the congruent signals (e.g., Adams et al., 2012; Adams & Kleck, 2003, 2005; Cushing et al., under review; Hadjikhani et al., 2008; Im et al., under review). Furthermore, in fearful faces this “pointing with the eyes” (Hadjikhani et al., 2008) to the source of threat disambiguates whence the threat is coming. When a fearful expression is combined with direct gaze, however, it tends to look less fearful due to the incongruity that direct gaze (an approach signal) creates in combination with the fearful expression (an avoidance signal), requiring more reflective processing to resolve the ambiguity inherent in the conflicting signal and the source of threat. Such interactions between gaze direction and a specific emotional facial expression (e.g., fear, joy, or anger) have been reported in many studies (e.g., Adams & Kleck, 2003, 2005; Adams et al., 2003; (Akechi et al., 2009; Bindemann, Burton, & Langton, 2008; Hess, Adams, & Kleck, 2007; Milders et al., 2011; Sander et al., 2007), suggesting that perceiving an emotional face involves integration of different types of social cues available in the face.

Recent findings suggest that visual threat stimuli may differentially engage the major visual streams – the magnocellular (M) and parvocellular (P) pathways. An emerging hypothesis posits that reflexive processing of clear threat cues may be predominantly associated with the more primitive, coarse, and action-oriented M pathway, while reflective, sustained processing of threat ambiguity may preferentially engage the slower, analysis-oriented P pathway (Adams et al., 2012; Adams & Kveraga, 2015; Kveraga & Bar, 2014). Indeed, recent fMRI studies directly compared M versus P pathway involvement in threat perception and supported this hypothesis by showing that congruent threat signals and incongruent threat signals in both face (Im et al., under review) and scene images (Kveraga, 2014) were processed preferentially by the M and P visual pathways, respectively. Moreover, Im et al. (under review) showed that observers’ trait anxiety levels differentially modulated M and P processing of clear and ambiguous facial threat cues such that higher anxiety facilitated processing of averted-gaze fear projected to M pathway and was associated with increased right amygdala reactivity, whereas higher anxiety impaired perception of direct-gaze fear projected to P pathway and was associated with increased left amygdala reactivity.

Another factor that is known to modulate emotional perception is sex-specific facial cues (Becker et al., 2007; Hess, Adams, & Kleck, 2004, 2005; Zebrowitz, Kikuchi, & Fellous, 2010). For example, anger is more readily perceived in male faces whereas joy is more readily perceived in female faces (Becker et al., 2007; Hess, Adams, & Kleck, 2004; Im et al., under review; see Adams, Hess, & Kleck, 2015 for review). Furthermore, the sex of the expresser also modulates the interaction between facial expression and gaze direction (Slepian et al., 2011). Despite the growing interests and empirical evidence in the role of sex as a biological variable in human behavioral and neuroimaging studies on emotional perception and behaviors, however, how observer’s sex plays a role in the differential attunements of M and P pathways to clear and ambiguous threat cues remains yet to be investigated.

Sex differences have been shown to be related to a variety of human brain and behavioral functions. In addition to cognitive differences including language (Shaywitz et al., 1995), navigational ability (Grön et al., 2000), defensiveness (Kline, Allen, & Schwartz, 1998), mathematical ability (Haier & Benbow, 1995), and attention (Mansour, Haier, & Buchsbaum, 1996), females and males also differ markedly in processing of affective stimuli (Larry Cahill, 2006; Campbell et al., 2002; Collignon et al., 2010; Hall, 1978; Hampson, van Anders, & Mullin, 2017; Stevens & Hamann, 2012). Behaviorally, females have been found to be more emotionally expressive than males (Kring & Gordon, 1998), possibly as a result of differences in socialization (Grossman & Wood, 1993). Moreover, females tend to be more emotionally reactive than males (Birnbaum & Croll, 1984; Shields, 1991) and tend to show stronger psychophysiological responses to emotional stimuli (Kring & Gordon, 1998; Orozco & Ehlers, 1998) and greater efficiency in using audio-visual, multisensory emotional information (Collignon et al., 2010) in order to recognize subtle facial emotions more accurately (Hoffmann et al., 2010) than males. The current study further aimed to investigate how observers’ sex affects perception and neural activity during the processing of compound threat cues from facial expression and eye gaze projected to M vs. P pathways.

One brain area that has been a frequent subject of sex differences research is the amygdala, a structure that is also known to play a critical role in the processing of affective information in general (Costafreda et al., 2008; Kober et al., 2008; Sergerie, Chochol, & Armony, 2008), and threat vigilance in particular (Davis & Whalen, 2001).

One of the consistent findings on the sex-related differences in amygdala activation is a different pattern of hemispheric lateralization. In male observers’ brains, the right amygdala was found to be dominant while in female observers’ brains, the left amygdala was found to be more involved in affective processing (Cahill et al., 1996; Cahill et al., 2001; Canli et al., 2000; Canli et al., 1999; Hamann et al., 1999; Killgore, Oki, & Yurgelun-Todd, 2001; Wager et al., 2003). Given that the amygdala’s involvement in facial expression and eye gaze interaction (Adams et al., 2012; Hadjikhani et al., 2008; Im et al., under review; Sato et al., 2004), as well as in processing each of them separately (Hoffman et al., 2007; Kawashima et al., 1999; Sergerie et al., 2008), has been well established, the current study was designed to focus on the functional and anatomical differences in the left and right amygdalae in female vs. male observers, during processing of face stimuli that convey different emotional expressions and gaze directions.

## Method

### Participants

One hundred and eight participants (64 females and 44 males) from the Massachusetts General Hospital (MGH) and surrounding communities participated in this study. Descriptive statistics for age, STAI-State and STAI-Trait are reported in Table 1 separately for males and females. The state and trait anxiety scores (STAI: Spielberger, 1983) of the female and male participants were not significantly different (STAI-state: *t*(106) = 0.925, *p* = 0.357; STAI-trait: *t*(106) = 0.364; *p* = 0.717). All had normal or corrected-to-normal visual acuity and normal color vision, as verified by the Snellen chart (Snellen, 1862), the Mars letter contrast sensitivity test (Arditi, 2005), and the Ishihara color plates (Ishihara, 1917). Informed consent was obtained from the participants in accordance with the Declaration of Helsinki. The experimental protocol was approved by the Institutional Review Board of MGH. The participants were compensated with $50 for their participation in this study.

**Table 1.** Participants’ descriptive statistics for age, STAI-State and STAI-Trait.

|  | **Mean Age** | **Mean STAI-S** | **Mean STAI-T** |
| --- | --- | --- | --- |
| **Female** | 36.39 (16.41) | 33.53 (9.92) | 35.3 (10.54) |
| **Male** | 37.69 (16.71) | 31.71 (8.45) | 32.55 (7.98) |

### Apparatus and stimuli

The stimuli were generated using MATLAB (Mathworks Inc., Natick, MA), together with the Psychophysics Toolbox extensions (Brainard, 1997; Pelli, 1997). The stimuli consisted of a face image presented in the center of a gray screen, subtending 5.79° x 6.78° of visual angle. We utilized a total of 24 face identities (12 female), 8 identities selected from the Pictures of Facial Affect (Ekman & Friesen, 1975), 8 identities from the NimStim Emotional Face Stimuli database (Tottenham et al., 2009), and the other 8 identities from the FACE database (Ebner, Riediger, & Lindenberger, 2010). The face images displayed either a neutral or fearful expression with either a direct gaze or averted gaze, and were presented as M-biased, P-biased, or Unbiased stimuli, resulting in 288 unique visual stimuli in total. Faces with an averted gaze had the eyes pointing either leftward or rightward.

Each face image was first converted to a two-tone image (black-white; termed the *Unbiased* stimuli from here on). From the two-tone image, low-luminance contrast (< 5% Weber contrast), achromatic, grayscale stimuli (M-biased stimuli), and chromatically defined, isoluminant red-green stimuli (P-biased stimuli) were generated. Examples of the Unbiased, M-biased, and P-biased stimuli are shown in Figure 1D. The low-luminance contrast images were designed to preferentially engage the M-pathway, while the isochromatic images were designed to engage the P-pathway, as such image manipulation has been employed successfully in previous studies (Awasthi et al., 2016; Cheng, Eysel, & Vidyasagar, 2004; Denison et al., 2014; Kveraga, Boshyan, & Bar, 2007; Schechter et al., 2003; Steinman, Steinman, & Lehmkuhle, 1997; Thomas et al., 2012). The foreground-background luminance contrast for achromatic M-biased stimuli and the isoluminance values for chromatic P-biased stimuli vary somewhat across individual observers. Therefore, these values were established for each participant in separate test sessions, with the participant positioned in the scanner, before commencing functional scanning. This ensured that the exact viewing conditions were subsequently used during functional scanning in the main experiment. Following the procedure in Kveraga et al. (2007), Thomas et al. (2012), and Boshyan et al. (in preparation), the overall stimulus brightness was kept lower for M stimuli (the average value of 115.88 on the scale of 0-255) than for P stimuli (146.06) to ensure that any processing advantages for M-biased stimuli were not due to greater overall brightness of the M stimuli, as described in detail below.

**Figure 1.**
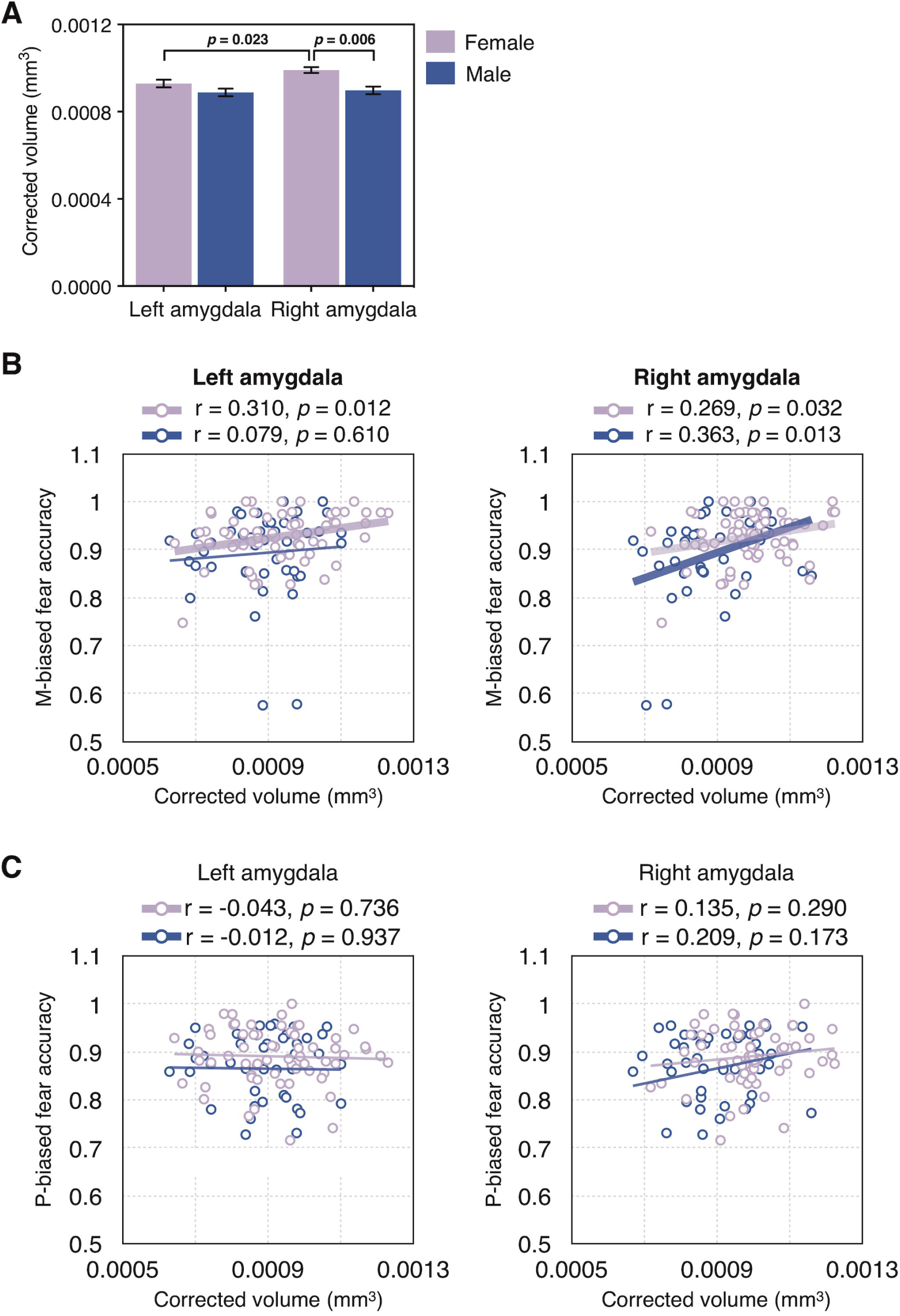
Sample trials of the pretests and the main experiment. **(A)** A sample trial of pretest 1 to measure the participants’ threshold for the foreground-background luminance contrast for achromatic M-biased stimuli. **(B)** A sample trial of pretest 2 to measure the participants’ threshold for the isoluminance values for chromatic P-biased stimuli. **(C)** A sample trial of the main experiment. **(D)** Sample images of unbiased, M-biased, and P-biased stimuli that were used in the main experiment.

### Procedure

Before the fMRI session, participants completed the Spielberger State-Trait Anxiety Inventory (STAI: Spielberger, 1983), followed by vision tests using the Snellen chart (Snellen, 1862), the Mars letter contrast sensitivity test (Arditi, 2005), and the Ishihara color plates (Ishihara, 1917). Participants were then positioned in the fMRI scanner and asked to complete the two pretests to identify the luminance values for M stimuli and chromatic values for P stimuli that were then used in the main experiment. The visual stimuli containing a face image were rear-projected onto a mirror attached to a 32-channel head coil in the fMRI scanner, located in a dimly lit room.

#### Pretest 1: Measuring luminance threshold for M-biased stimuli

The appropriate luminance contrast was determined by finding the luminance threshold via a multiple staircase procedure. Figure 1A illustrates a sample trial of Pretest 1. Participants were presented with visual stimuli for 500 msec and instructed to make a key press to indicate the facial expression of the face that had been presented. They were required to choose one of the four options: 1) neutral, 2) angry, 3) fearful, or 4) did not recognize the image. One-fourth of the trials were catch trials in which the stimulus did not appear. To find the threshold for foreground-background luminance contrast, our algorithm computed the mean of the turnaround points above and below the gray background ([120 120 120] RGB value on the 8-bit scale of 0-255). From this threshold, the appropriate luminance (∼3.5% Weber contrast) value was computed for the face images to be used in the low-luminance-contrast (M-biased) condition. As a result, the average foreground RGB values for M-biased stimuli were [116.71 116.71 116.71] ± 2.02 (SD) for female participants and [116.08 116.08 116.08] ± 2.19 (SD) for male participants.

#### Pretest 2: Measuring red-green isoluminance value for P-biased stimuli

Figure 1B illustrates a sample trial of Pretest 2. For the chromatically defined, isoluminant (P-biased) stimuli, each participant’s isoluminance point was determined using heterochromatic flicker photometry with two-tone face images displayed in rapidly alternating colors, between red and green. The alternation frequency was ∼14Hz, because in our previous studies (Boshyan et al., in preparation; Kveraga et al., 2007; Kveraga & Bar, 2014; Thomas et al., 2012) we obtained the best estimates for the isoluminance point (e.g., narrow range within-subjects and low variability between-subjects; Kveraga et al., 2007) at this frequency. The isoluminance point was defined as the color values at which the flicker caused by luminance differences between red and green colors disappeared and the two alternating colors fused, making the image look steady. On each trial, participants were required to report via a key press whether the stimulus appeared flickering or steady. Depending on the participant’s response, the value of the red gun in [r g b] was adjusted up or down in a pseudorandom manner for the next cycle. The average of the values in the narrow range when a participant reported a steady stimulus became the isoluminance value for the subject used in the experiment. Thus, isoluminant stimuli were defined only by chromatic contrast between foreground and background, which appeared equally bright to the observer. On the background with green value of 140, the resulting foreground red value was 151.34 ± 3.86 (SD) on average for female participants and 150.79 ± 4.33 (SD) on average for male participants. Therefore, the isoluminant P-biased stimuli were objectively brighter than the low-luminance contrast, M-biased, stimuli. This was done to ensure that any performance advantages for the M-biased stimuli over the P-biased stimuli (as found in Kveraga et al., 2007) were due to pathway-biasing and not stimulus brightness.

#### Main experiment

Figure 1C illustrates a sample trial of the main experiment. After a variable pre-stimulus fixation period (200-400 msec), a face stimulus was presented for 1000 msec, followed by a blank screen (1100-1300 msec). Participants were required to indicate whether a face image looked fearful or neutral, as quickly as possible. Key-target mapping was counterbalanced across participants: One half of the participants pressed the left key for neutral and the right key for fearful and the other half pressed the left key for fearful and the right key for neutral. Feedback was provided on every trial. The accuracy (proportion correct) of participants’ responses and the response time (RT) were recorded and analyzed as behavioral measurement.

#### fMRI data acquisition and analysis

fMRI images of brain activity were acquired using a 1.5 T scanner (Siemens Avanto) with a 32-channel head coil. High-resolution anatomical MRI data were acquired using T1-weighted images for the reconstruction of each subject’s cortical surface (TR=2300 ms, TE=2.28 ms, flip angle=8°, FoV=256x256 mm^^2^^, slice thickness=1 mm, sagittal orientation). The functional scans were acquired using simultaneous multislice, gradient-echo echoplanar imaging with a TR of 2500 ms, three echoes with TEs of 15 ms, 33.83 ms, and 52.66 ms, flip angle of 90°, and 58 interleaved slices (3x3x2 mm resolution). Scanning parameters were optimized by manual shimming of the gradients to fit the brain anatomy of each subject, and tilting the slice prescription anteriorly 20-30° up from the AC-PC line as described in the previous studies (Deichmann et al., 2003; Kveraga et al., 2007; Wall, Walker, & Smith, 2009), to improve signal and minimize susceptibility artifacts in the subcortical brain regions. For each participant, the first 15 seconds of each run were discarded, followed by acquisition of 96 functional volumes per run (lasting 4 minutes). There were four successive functional runs, providing the 384 functional volumes per subject in total, including the 96 null, fixation trials and the 288 stimulus trials. In our 2 (Emotion: Fear and Neutral) x 2 (Eye gaze direction: Direct gaze vs. Averted gaze) x 3 (Bias: Unbiased, M-biased, and P-biased) design, each condition had 24 trials, and the sequence of total 384 trials was optimized for hemodynamic response estimation efficiency using the *optseq2* software (https://surfer.nmr.mgh.harvard.edu/optseq/).

The acquired functional images were pre-processed using SPM8 (Wellcome Department of Cognitive Neurology). The functional images were corrected for differences in slice timing, realigned, corrected for movement-related artifacts, coregistered with each participant’s anatomical data, normalized to the Montreal Neurological Institute (MNI) template, and spatially smoothed using an isotropic 8-mm full width half-maximum (FWHM) Gaussian kernel. Outliers due to movement or signal from preprocessed files, based on thresholds of 3 SD from the mean, 0.75 mm for translation and 0.02 radians rotation, were removed from the data sets, using the ArtRepair software (Mazaika et al., 2009).

#### Whole brain analysis

For whole brain analyses, subject-specific contrasts were estimated using a fixed-effects model. These contrast images were used to obtain subject-specific estimates for each effect then entered into a second-level analysis treating participants as a random effect, using one-sample t-tests at each voxel. Age and anxiety of participants were controlled as covariates. One-sample t-tests were first conducted across all subjects for each of the conditions of our interest, compared to the baseline (Null trials). In order to examine the sex difference in the pattern of whole brain activation, we then conducted two-sample t-tests between female and male participants for each condition. For illustration purposes, the resulting t-test images showing the difference between female and male participants were overlaid onto a group average brain of the 108 participants, using the Multi-image Analysis GUI (Mango: http://rii.uthscsa.edu/mango/index.html) software. For visualization of the contrasts (Figure 3), we used the threshold of *p* < 0.001 (uncorrected) with a minimal cluster size of 5 voxels. These parameters are more conservative than those that have been argued to optimally balance between Type 1 and Type 2 errors (Lieberman & Cunningham, 2009; height *p* < 0.005, uncorrected, extent: 10 voxels, see also Adams et al., 2012; Kveraga et al., 2011). To report all the significant clusters in Table 2, we used the threshold of *p* < 0.05, FWE whole-brain corrected.

**Table 2.** BOLD activations from group analysis, thresholded at *p* < 0.05, FWE corrected. **– indicates that this cluster is part of a larger cluster immediately above.** **\* indicates p < 0.005, uncorrected.**

| **M-biased direct fear** | Extent | Region label | t | x | y | z |
| --- | --- | --- | --- | --- | --- | --- |
| Female > Male | 3956 | L Superior occipital gyrus | 4.422 | -33 | -85 | 30 |
|  | 3956 | R Precuneus | 3.747 | 9 | -64 | 56 |
|  | 3956 | L Cuneus | 3.562 | -6 | -82 | 32 |
|  | - | R Supplementary motor area | 2.702 | 9 | -22 | 62 |
|  | - | L Supplementary motor area | 2.909 | -6 | 8 | 68 |
|  | - | L Anterior cingulate cortex | 2.396 | -6 | 20 | 44 |
|  | - | R Anterior cingulate cortex | 3.162 | 6 | 26 | 34 |
|  | - | R Superior frontal gyrus | 2.76 | 18 | 38 | 48 |
| Male > Female |  | None |  |  |  |  |
| **M-biased averted fear** | Extent | Region label | t | x | y | z |
| Female > Male |  | None |  |  |  |  |
| Male > Female | 3188 | R Parahippocampal cortex | 3.879 | 27 | -28 | -4 |
|  | - | R Amygdala | 3.193 | 24 | -7 | -16 |
|  | 3188 | R Cerebellum | 3.531 | 27 | -67 | -20 |
|  | 3188 | L Parahippocampal cortex | 2.369 | -30 | -40 | -8 |
|  | 3188 | R Retrosplenial cortex | 2.029 | 18 | -61 | 18 |
|  | 3188 | R Hippocampus | 2.801 | 27 | -28 | -8 |
|  | 1621 | L Middle temporal gyrus | 2.668 | -48 | -22 | -16 |
| **P-biased direct fear** | Extent | Region label | t | x | y | z |
| Female > Male | 4682 | L Anterior cingulate cortex | 2.633 | -12 | 41 | 14 |
|  | 4682 | R Anterior cingulate cortex | 3.574 | 3 | 53 | 12 |
|  | - | L Medial prefrontal cortex | 2.243 | -12 | 53 | 8 |
|  | 4682 | L Insula | 4.014 | -42 | -7 | 8 |
|  | - | L Superior temporal sulcus | 3.011 | -45 | -31 | 4 |
|  |  |  | 3.149 | -45 | -61 | 14 |
|  | 4682 | R Superior temporal sulcus | 2.132 | 63 | -43 | 10 |
|  | 4682 | L Dorsolateral prefrontal | 3.735 | -42 | 38 | 12 |
|  | 4682 | R Fusiform cortex | 3.261 | 51 | -55 | -24 |
| Male > Female |  | None |  |  |  |  |
| **P-biased averted fear** | Extent | Region label | t | x | y | z |
| Female > Male |  | None |  |  |  |  |
| Male > Female |  | None |  |  |  |  |

| **M-biased direct neutral** | Extent | Region label | t | x | y | z |
| --- | --- | --- | --- | --- | --- | --- |
| Female > Male | 3643 | L Fusiform cortex | 2.95 | -45 | -55 | -26 |
|  | 3643 | L Superior temporal gyrus | 2.341 | -51 | -46 | 22 |
|  | - | L Intraparietal sulcus | 2.621 | -51 | -40 | 38 |
|  | - | L Supramarginal cortex | 3.125 | -51 | -25 | 22 |
|  | - | L Superior temporal gyrus | 2.259 | -51 | -25 | 12 |
|  | 3643 | Middle cingulate cortex | 3.632 | 0 | 17 | 36 |
|  | - | Supplementary motor area | 2.7 | 0 | 20 | 48 |
|  | 3643 | L Precentral gyrus | 2.36 | -27 | -13 | 70 |
|  | 3643 | L Insula | 2.055 | -39 | -10 | 10 |
| Male > Female |  | None |  |  |  |  |
| **M-biased averted neutral** | Extent | Region label | t | x | y | z |
| Female > Male | 2040 | L Amygdala | 2.918 | -30 | -4 | -14 |
|  | 2040 | R Hippocampus | 1.895 | 36 | -22 | -12 |
|  | 2040 | R Temporal pole | 3.77 | 36 | 20 | -26 |
|  | 1435 | R Fusiform cortex | 2.647 | 48 | -34 | -26 |
|  | - |  | 3.572 | 48 | -49 | -24 |
|  | 2040 | L Putamen | 3.351 | -30 | -1 | -8 |
|  | 2040 | L Orbitofrontal cortex | 2.046 | -27 | 23 | -10 |
|  | 4628 | L Precuneus | 2.957 | -18 | -52 | 58 |
|  | 4628 | R Anterior cingulate cortex | 2.955 | 9 | 41 | 20 |
|  | 4628 | R Precuneus | 3.372 | 9 | -67 | 58 |
|  | 2040 | R Inferior temporal gyrus | 3.377 | 36 | -4 | -36 |
|  | 4628 | L Superior occipital gyrus | 4.098 | -42 | -73 | 16 |
|  | - | L Superior temporal sulcus | 3.613 | -45 | -64 | 16 |
|  | 4628 | L Postcentral gyrus | 2.616 | -45 | -19 | 56 |
|  | 2040 | L Medial orbitofrontal cortex | 2.818 | -18 | 14 | -18 |
|  |  |  | 2.898 | -3 | 20 | -18 |
|  | 4628 | L Superior frontal gyrus | 3.475 | -3 | 29 | 52 |
|  |  |  | 3.331 | -3 | 62 | 38 |
| Male > Female |  | None |  |  |  |  |
| **P-biased direct neutral** | Extent | Region label | t | x | y | z |
| Female > Male | 4476 | L Pallidum | 3.936 | -21 | -10 | -4 |
|  | 4476 | L Parahippocampal cortex | 2.147 | -21 | -40 | -4 |
|  | 4476 | L Amygdala | 1.66 | -21 | -7 | -12 |
|  | 4476 | L Superior temporal sulcus | 3.888 | -63 | -49 | 0 |
|  | - | L Superior temporal gyrus | 2.562 | -63 | -31 | 4 |
|  | 4476 | L Insula | 2.575 | -39 | -19 | 8 |
|  | 4476 | L Inferior frontal gyrus | 2.128 | -39 | 35 | 12 |
|  | 4476 | L Fusiform cortex | 2.943 | -54 | -49 | -22 |
|  | - | L Middle temporal gyrus | 1.823 | -54 | -7 | -26 |
|  | 4476 | L Supramarginal gyrus | 3.731 | -36 | -55 | 20 |
| Male > Female |  | None |  |  |  |  |
| **P-biased averted neutral** | Extent | Region label | t | x | y | z |
| Female > Male | 4 | L Amygdala* | 3.359 | -21 | -1 | -12 |
| Male > Female | 2066 | R Posterior cingulate cortex | 4.129 | 3 | -52 | 16 |
|  | 2066 | R Precuneus | 3.376 | 3 | -46 | 50 |
|  | 2066 | R Cuneus | 3.14 | 12 | -70 | 38 |
|  | 1517 | R Middle frontal gyrus | 3.477 | 42 | 23 | 42 |
|  | 1517 | R Insula | 3.004 | 42 | 5 | 12 |
|  | 1517 | R Precentral gyrus | 2.78 | 48 | -13 | 52 |
|  | 1517 | R Inferior frontal gyrus | 2.636 | 48 | 20 | 6 |
|  | 2066 | R Thalamus | 2.755 | 15 | -19 | 0 |
|  | 1517 | R Postcentral gyrus | 2.331 | 54 | -22 | 54 |

#### Region-of-Interest (ROI) analysis

For ROI analyses, we used the rfxplot toolbox (http://rfxplot.sourceforge.net) for SPM and extracted the beta weights from the left and right amygdala. The coordinates of the left and right amygdala (x=±20, y=−4, z=-15) were defined by functionally restricting brain activation based on the contrast of Unbiased fear minus Unbiased neutral, collapsed across the eye gazes using random effects models (height: *p* < 0.01, uncorrected; extent: 5 voxels), within the anatomical label for the left and right amygdala (obtained by the anatomical parcellation of the normalized brain; Tzourio-Mazoyer et al., 2002). Around these coordinates, we defined 6mm spheres and extracted all the voxels from each individual participant’s functional data within those spheres. The extracted beta weights were subjected to mixed repeated measures ANOVA, conducted separately for Fearful and Neutral face stimuli.

#### Estimation of Amygdala volume

To assess amygdala volumes, we performed quantitative morphometric analysis of T1– weighted MRI data using an automated segmentation and probabilistic ROI labeling technique (FreeSurfer, http://surfer.nmr.mgh.harvard.edu). This procedure has been widely used in volumetric studies and was shown to be comparable in accuracy to that of manual labeling (Bickart et al., 2011; Fischl et al., 2002). The estimated amygdala volumes for each individual participant were divided by total intracranial volume of the participant in order to adjust for individual differences in head size, as performed in the prior work (e.g., OLJBrien et al., 2006)

## Results

### Behavioral results: accuracy

Figures 2A and 2B show the average accuracy (proportion correct) of female and male participants, separately plotted for fearful face stimuli and neutral face stimuli, respectively. For behavioral responses to fearful faces, a mixed repeated measures ANOVA with Sex (Female and Male) as a between-subject factor (our main interest) and with Bias (2 levels: M-biased and P-biased) and Eye gaze (2 levels: Direct gaze and Averted gaze) as within-subject factors revealed three main effects. First, there was a significant main effect of Sex (*F*(1,106) = 5.658, *p* = 0.019) with female participants being more accurate than male participants for fear stimuli overall. Second, there was also a significant main effect of Bias (*F*(1,106) = 28.81, *p* < 0.001) with M-biased fear stimuli being recognized more accurately than P-biased fear stimuli. Third, there was a significant main effect of Eye gaze (*F*(1,106) = 57.39, *p* < 0.001), with averted gaze fear expressions being recognized more accurately than direct eye gaze fear expressions. Neither the two-way interactions between the factors nor the three-way interaction of all the factors was significant (*p* > 0.481). Greater accuracy for averted fear than direct fear faces overall was also observed in the previous studies showing that fearful faces with averted eye gaze tended to be perceived as more intense compared to fearful faces with direct eye gaze (e.g., Adams & Kleck, 2005). Moreover, greater accuracy for M-biased fear than P-biased fear faces has also been observed in our recent work (Cushing et al., under review). Since our main interest was to test sex differences in perception of emotional face with different eye gaze directions and pathway biases, we further conducted planned comparisons between female and male participants for each condition, and found that the M-biased averted fear (*t*(106) = 2.76, *p* = 0.007) and P-biased averted fear (*t*(106) = 1.757, *p* = 0.08, marginally significant) yielded significantly higher accuracy for female than male participants.

**Figure 2.**
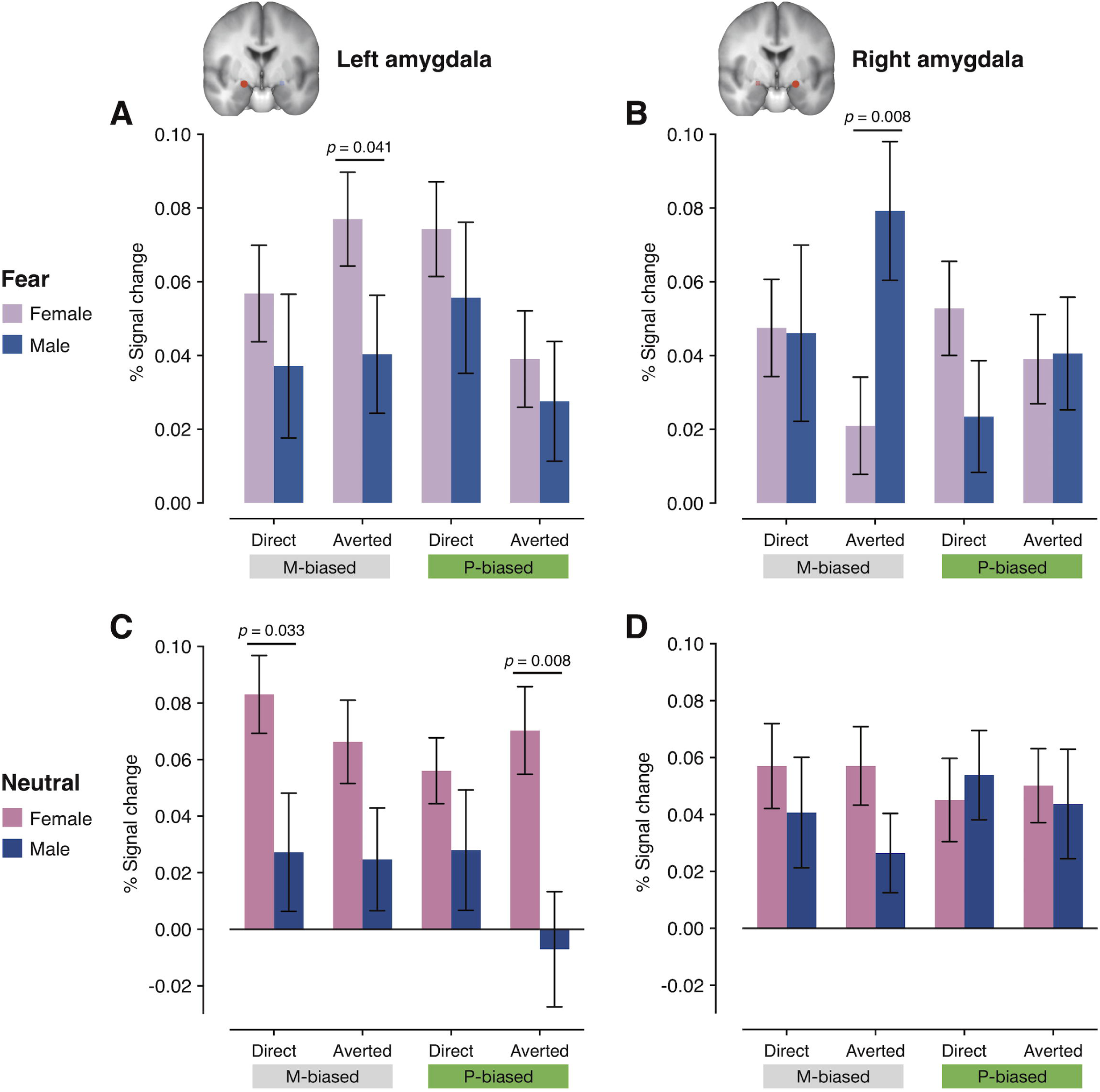
Behavioral results. **(A)** The mean accuracy for fearful faces paired with direct or averted eye gazes, presented in M-or P-biased stimuli. The pink bar graphs indicate the accuracy for female participants and the blue bar graphs indicate the accuracy for male participants. The error bars indicate the standard error of the mean (SEM). **(B)** The median RT for fearful faces paired with direct or averted eye gazes, presented in M-or P-biased stimuli. The pink bar graphs indicate the RT for female participants and the blue bar graphs indicate the RT for male participants. The error bars indicate the standard error of the mean (SEM). **(C)** The mean accuracy for neutral faces paired with direct or averted eye gazes, presented in M-or P-biased stimuli. The pink bar graphs indicate the accuracy for female participants and the blue bar graphs indicate the accuracy for male participants. The error bars indicate the standard error of the mean (SEM). **(D)** The median RT for neutral faces paired with direct or averted eye gazes, presented in M-or P-biased stimuli. The pink bar graphs indicate the RT for female participants and the blue bar graphs indicate the RT for male participants. The error bars indicate the standard error of the mean (SEM).

For neutral face stimuli, mixed repeated measures ANOVA with Sex (Female and Male) as a between-subject factor (our main interest) and with Bias (2 levels: M-biased and P-biased) and Eye gaze (2 levels: Direct gaze and Averted gaze) as within-subject factors showed only marginally significant main effect of Sex with female participants being more accurate than male participants (*F*(1,106) = 3.618, *p* = 0.060), significant main effect of Bias (*F*(1,106) = 8.181, *p* = 0.005) with M-biased stimuli being recognized more accurately than P-biased stimuli, and significant main effect of Eye gaze (*F*(1,106) = 8.306, *p* = 0.005) with averted eye gaze being recognized more accurately than direct eye gaze. Although the two-way interaction of Bias x Sex (*F*(1,106) = 0.006, *p* = 0.938) and interaction of Bias x Eye gaze (*F*(1,106) = 0.459, *p* = 0.50) were not significant, the two-way interaction of Eye gaze x Sex was marginally significant (*F*(1,106) = 3.743, *p* = 0.056). We assessed the nature of the Eye gaze x Sex interaction by using post hoc pairwise comparisons (Bonferroni corrected) and found that female participants recognized neutral faces with averted eye gaze more accurately than with direct eye gaze (*p* = 0.024 for M-biased and *p* = 0.015 for P-biased), while male participants did not show any difference in the accuracy for neutral faces with direct eye gaze vs. averted eye gaze (*p*’s > 0.841). The three-way interaction among the factors was not significant (*F*(1,106) = 0.815, *p* = 0.369). In order to test the sex difference in the accuracy for each of the conditions, we also conducted further planned comparisons between female and male participants for each of the four conditions, and found that female participants were more accurate than male participants at recognizing neutral faces with averted gaze, both in M-biased (*p* = 0.031) and in P-biased (*p* = 0.017) stimuli.

#### Behavioral results: response time (RT)

Figure 2C shows the median RT of female and male participants for fearful face stimuli. Only the RTs for the correct trials were used for the analyses, and outliers (3SD above the group mean) within each condition were screened. As a result, 1.03% of the data points on average were excluded for the further analyses. The mixed repeated measures ANOVA with Sex (Female and Male) as a between-subject factor (our main interest) and with Bias (2 levels: M-biased and P-biased) and Eye gaze (2 levels: Direct gaze and Averted gaze) as within-subject factors showed no significant main effects of Sex (*F*(1,106) = 0.014, *p* = 0.906) or Bias (*F*(1,106) = 0.960, *p* = 0.329), although a main effect of Eye gaze was significant with direct-gaze fear faces recognized faster than averted-gaze fear faces (*F*(1,106) = 38.142, *p* < 0.001). We also found that the two-way interaction of Eye gaze x Sex (*F*(1,106) = 3.671, *p* = 0.058, marginally significant) and the interaction of Bias and Eye gaze were significant (*F*(1,106) = 7.029, *p* = 0.009). In order to assess the nature of the significant two-way interactions, we conducted post hoc pairwise comparisons (Bonferroni corrected) and found that male participants were significantly faster for P-biased fearful faces with direct gaze than with averted gaze (*p* = 0.027) although female participants did not show significant differences in RT between direct vs. averted fear. Moreover, P-biased fearful faces were recognized significantly faster with direct gaze than for those with averted gaze (*p* = 0.031), while M-biased fearful faces did not show any significant eye gaze effects (*p* > 0.761). Thus, the significant main effect of Eye gaze we found seemed to be driven mainly by faster RT for P-biased gaze fear faces. The three-way interaction of all the factors was not significant (*F*(1,106) = 0.047, *p* = 0.828).

Figure 2D shows the RT of female and male participants for neutral face stimuli. The mixed repeated measures ANOVA with Sex (Female and Male) as a between-subject factor (our main interest) and with Bias (2 levels: M-biased and P-biased) and Eye gaze (2 levels: Direct gaze and Averted gaze) as within-subject factors showed that the main effect of Sex was not significant (*F*(1,106) = 0.009, *p* = 0.926), although the main effects of Bias and Eye gaze were significant, with M-biased neutral faces being recognized faster than P-biased neutral faces (*F*(1,106) = 6.842, *p* = 0.010) and neutral faces with direct gaze being recognized faster than with averted gaze (*F*(1,106) = 8.814, *p* = 0.004). None of the two-way or three-way interaction was significant (*p* > 0.221). Together, we found these sex differences in the perception of faces: Compared to male, female observers showed more accurate perception of both fearful and neutral faces with averted gaze, suggesting that they are more sensitive to covert information conveyed by a non-emotional facial cue, such as eye gaze.

#### fMRI results

Figure 3 shows different patterns of left and right amygdala activations in female and male participants when they viewed fear and neutral faces with direct and averted eye gazes in M- and P-biased stimuli. For M-biased fearful faces with averted eye gaze (Figure 3A), female participants showed greater left amygdala activation (dorsal amygdala/SI, peak: [x=-24, y=−7, z= −11]) than male participants, whereas male participants showed greater right amygdala activation (peak: [x=24, y=−7, z=−16]) than female participants. For P-biased fearful faces with direct eye gaze, female participants showed greater left amygdala activation (dorsal amygdala/SI, peak: [x=-27, y=−2, z=-12]). For neutral faces, female participants showed greater left dorsal amygdala activation than male participants for M-biased averted gaze (peak: [x=-30, y=−4, z=-14])), P-biased direct gaze (peak: [x=-21, y=−7, z=-12]), and P-biased averted gaze (peak: [x=-21, y=−1, z=-12]) conditions. These results suggest that compound cues of emotional expression and eye gaze direction of face stimuli projected to M- and P-pathways are processed differentially in the left and right amygdala of female vs. male observers. The complete list of the brain areas that showed significant differences between female vs. male participants for each contrast is shown in Table 2.

**Figure 3.**
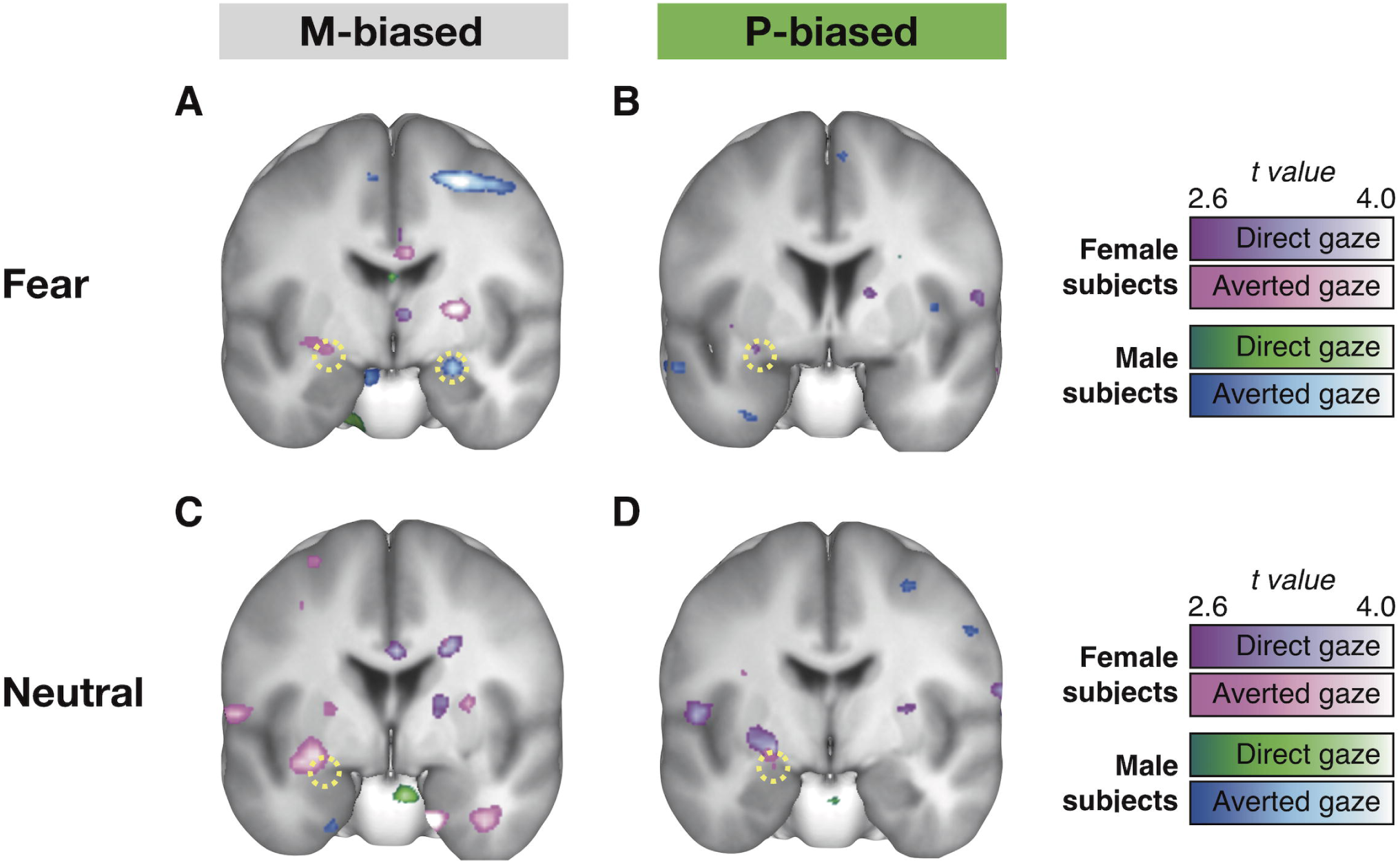
Different patterns of left and right amygdala activations in female and male participants when they viewed: **(A)** M-biased fearful faces, **(B)** P-biased fearful faces, **(C)** M-biased neutral faces, and **(D)** P-biased neutral faces.

Because the whole brain analyses between female vs. male participants revealed significant differences in left and right amygdala activations, we next conducted regions of interest (ROI) analysis to directly compare the amygdala responsivity to different combinations of emotional expression, eye gaze direction, and pathway bias in female vs. male participants. The coordinates of the left and right amygdala were determined from an independent contrast of Unbiased fearful vs. Unbiased neutral face stimuli (x=±20, y=−4, z=-15). Figure 4A shows the % signal change in the left amygdala for four different fear conditions (2 bias: M- and P-by 2 eye gaze directions: direct and averted), plotted separately for female and male participants, next to each other. The mixed repeated measures ANOVA with Sex (Female and Male) as a between-subject factor (our main interest) and with Bias (2 levels: M-biased and P-biased) and Eye gaze (2 levels: Direct gaze and Averted gaze) as within-subject factors showed a significant main effect of Sex (*F*(1,106) = 3.945, *p* = 0.049) with female participants showing greater left amygdala activation than male participants. Neither the main effect of Bias (*F*(1,106) = 0.149, *p* = 0.700) nor the main effect of Eye gaze (*F*(1,106) = 0.789, *p* = 0.376) was significant. We also found a significant two-way interaction between Bias and Eye gaze (*F*(1,106) = 5.011, *p* = 0.027), although the other two-way or three-way interactions were not significant (*p* > 0.492). The nature of the significant interaction between Bias and Eye gaze was further tested for female and male participants separately using the Bonferroni pairwise comparison. We found that the left amygdala reactivity was significantly higher for P-biased direct fear than for P-biased averted fear (*p* = 0.045) in female participants, but not in male participants (*p* > 0.583). Finally, we tested the sex difference by using further planned comparisons between female and male participants for each of the four conditions, and found that female participants showed significantly greater left amygdala activation for the M-biased averted fear than male participants (*p* = 0.041), although the other conditions did not reach significant difference between sex groups (*p*’s > 0.385).

**Figure 4.**
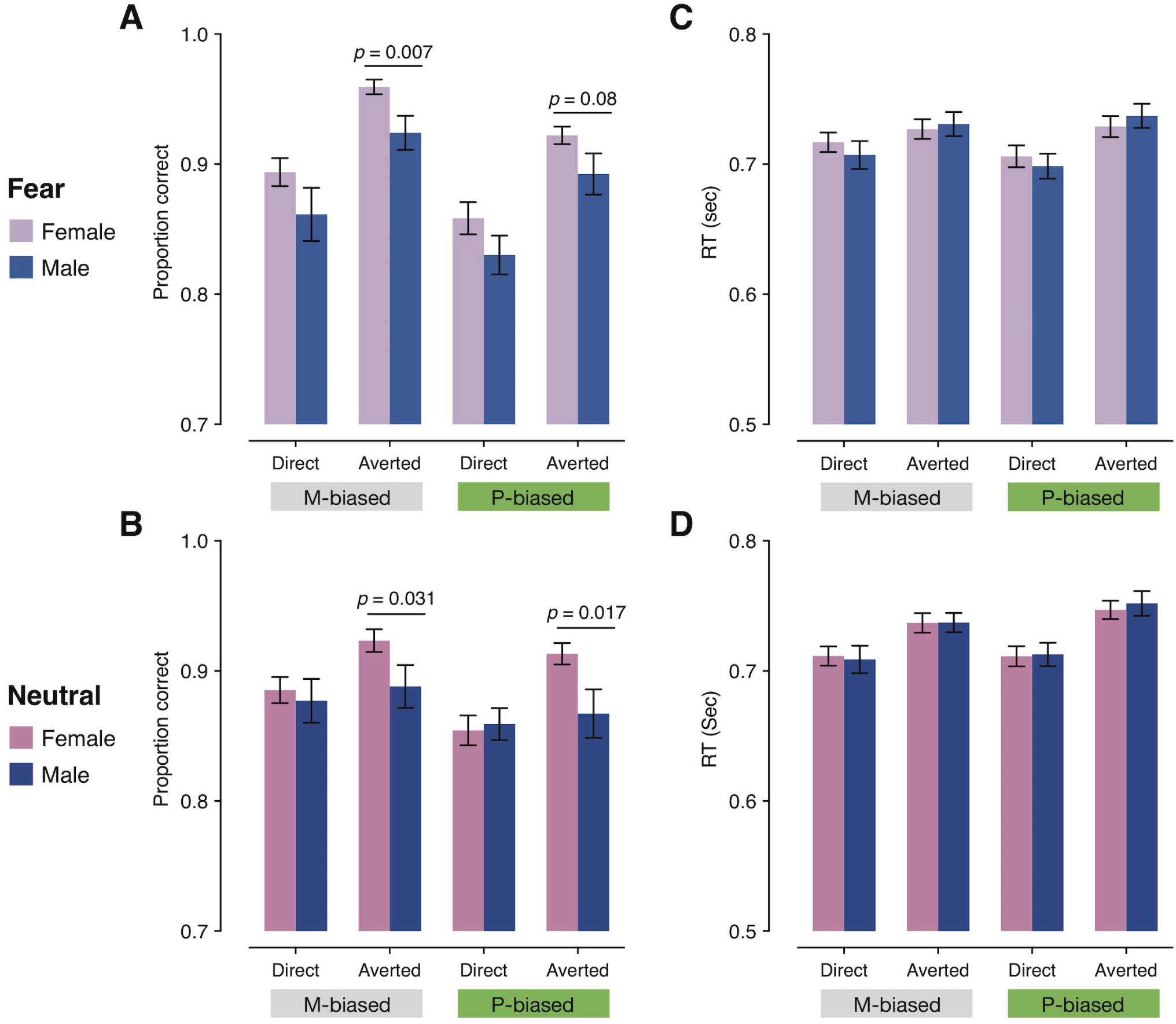
The % signal change resulting from the ROI analyses. **(A)** The % signal change of the left amygdala for fearful faces with direct or averted eye gaze in M-biased or P-biased stimuli. **(B)** The % signal change of the right amygdala for fearful faces with direct or averted eye gaze in M-biased or P-biased stimuli. **(C)** The % signal change of the left amygdala for neutral faces with direct or averted eye gaze in M-biased or P-biased stimuli. **(D)** The % signal change of the right amygdala for neutral faces with direct or averted eye gaze in M-biased or P-biased stimuli.

Figure 4B shows the right amygdala activation. The mixed repeated measures ANOVA with Sex (Female and Male) as a between-subject factor (our main interest) and with Bias (2 levels: M-biased and P-biased) and Eye gaze (2 levels: Direct gaze and Averted gaze) as within-subject factors showed no statistically significant main effects of Sex, Bias, or Eye gaze (*p*’s > 0.318). Two-way interactions between Bias and Sex (*F*(1,106) = 5.026, *p* = 0.027) and between Eye gaze and Sex (*F*(1,106) = 5.985, *p* = 0.016), but not between Bias and Eye gaze (*F*(1,106) = 0.006, *p* = 0.939), were significant. Further contrast analyses conducted separately for female and male participants revealed that the right amygdala activation was greater for M-biased than P-biased stimuli in male participants (*p* = 0.046), but slightly greater for P-biased than M-biased stimuli in female participants (although the trend did not reach significance: *p* = 0.281). Moreover, the right amygdala activation was greater for averted fear faces both in M- and P-biased stimuli in male participants (*p* = 0.048), but slightly greater for direct fear faces both in M- and P-biased stimuli in female participants (*p* = 0.052). Male participants also showed significantly greater right amygdala activation for M-biased averted fear face than any of the other three conditions (using the contrast weight of [−1 +3 −1 −1], *p* = 0.041). This is consistent with the previous findings suggesting that right amygdala is sensitive to detecting congruent, clear threat cues (Adams et al., 2012; Cushing et al., under review; Cushing et al., under review; Im et al., under review) particularly via M-biased projection (Cushing et al., under review; Im et al., under review). This pattern, however, was not significant in female participants. Finally, further planned comparisons between female and male participants for each of the four conditions showed significantly greater right amygdala activation for M-averted fear in male participants than in female participants (*p* = 0.008).

The left amygdala showed greater activation for neutral face stimuli overall in female participants than in male participants (Figure 4C). A mixed repeated measures ANOVA with Sex (Female and Male) as a between-subject factor (our main interest) and with Bias (2 levels: M-biased and P-biased) and Eye gaze (2 levels: Direct gaze and Averted gaze) as within-subject factors showed a significant main effect of Sex (*F*(1,106) = 9.829, *p* = 0.002), supporting this observation. None of the other main effects or interactions were significant (*p*’s > 0.200). Planned comparisons between female and male participants for each of the four conditions showed significantly greater left amygdala activation for M-biased direct neutral (*p* = 0.033) and P-biased averted neutral (*p* = 0.008) in female participants than in male participants. For the right amygdala responses to neutral face stimuli (Figure 4D), however, none of the main effects or the interactions was significant (*p*’s > 0.201). Therefore, the fMRI results suggest that male participants show greater right amygdala activation for M-biased and averted fear face stimuli (e.g., clear threat cue; Adams et al., 2012; Cushing et al., under review; Im et al., under review) indicating greater attunement of the right amygdala to clear threat conveyed by the magnocellular pathway, whereas female participants show greater left amygdala involvement in processing of faces (both fearful and neutral), compared to male participants.

In addition to the differences in the BOLD responses, we also found differences in the volume (divided by total intracranial volume to correct for variable head size) of the left and right amygdala between female and male participants (Figure 5A). Mixed repeated measures ANOVA with Sex (Female and Male) as a between-subject factor (our main interest) and with Hemisphere (2 levels: Left and Right) as a within-subject factors showed a significant main effect of Sex (*F*(1,106) = 9.519, *p* = 0.003) with the amygdala volume being greater in female than male participants, a significant main effect of the Hemisphere (*F*(1,106) = 13.119, *p* < 0.001) with the right amygdala volume being greater than the left amygdala volume, and a significant interaction (*F*(1,106) = 7.161, *p* = 0.009). Further t-tests (Bonferroni corrected) revealed the nature of the interaction: in female participants the amygdala volume was significantly greater in the right than in the left (*p* = 0.023), whereas male participants did not show a significant difference between the volumes of the left and right amygdala (*p* = 0.988); lastly, the right amygdala volume was significantly greater in female participants than in male participants (*p* = 0.006).

**Figure 5.**
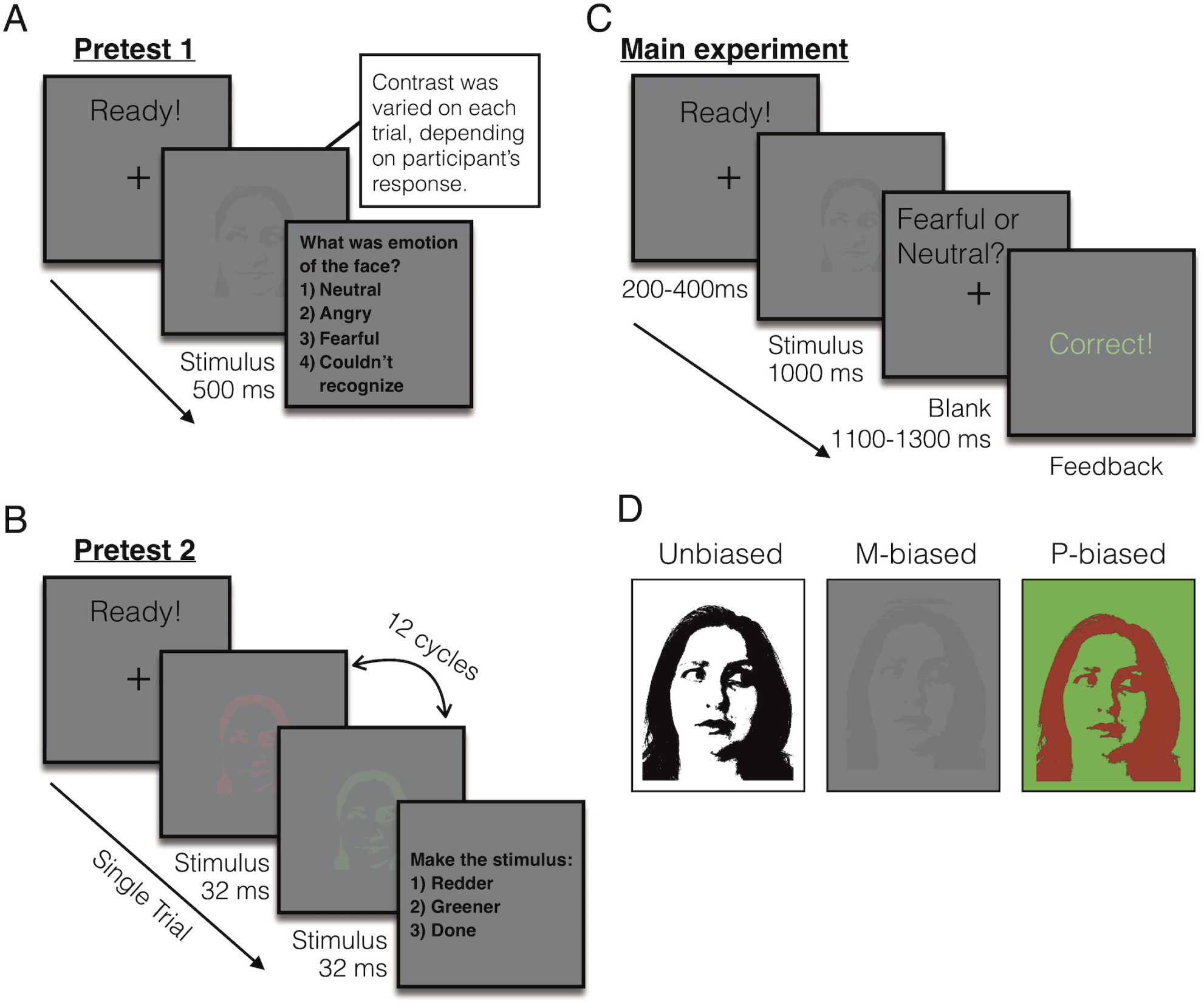
The results of the analyses of amygdala volumes. **(A)** The volume of the left and right amygdala for female and male participants. **(B)** The correlation between the left and the right amygdala volume and the behavioral accuracy for M-biased fear. The pink dots indicate female participants and the blue dots indicate male participants. The thinker regression lines indicate the statistically significant correlations. **(C)** The correlation between the left and the right amygdala volume and the behavioral accuracy for P-biased fear. The pink dots indicate female participants and the blue dots indicate male participants.

Finally, the measures of the left and right amygdala volume were found to correlate with observers’ behavioral accuracy for fearful faces differentially in female vs. male participants. In female participants (pink dots and regression lines in Figure 5B), both the left and right amygdala volumes showed significant positive correlation with their accuracy for M-biased fear (averaged across direct and averted eye gaze), with slightly stronger correlation for the left amygdala than for the right amygdala (left amygdala: *r* = 0.310, *p* = 0.012; right amygdala: *r* = 0.269, *p* = 0.032). In male participants (blue dots and regression lines in Figure 5B), however, only the right amygdala showed significant correlation with their behavioral accuracy for M-biased fear (*r* = 0.363, *p* = 0.013), but not the left amygdala (*r* = 0.079, *p* = 0.610). Unlike the accuracy for M-biased fear, neither the accuracy for P-biased fear faces (Figure 5C) nor M- and P-biased neutral faces showed correlation with the amygdala volumes (all *p’s* > 0.173). Together, our findings suggest that female and male participants show differences in perception of emotion from compound cues of facial expressions and eye gaze directions and in hemispheric lateralization of amygdala responsivity and volumes.

## Discussion

The goal of this study was to explore behavioral and neural differences between female and male observers during perception of faces with potential threat cues conveyed by the combination of facial expression (fearful or neutral) and eye gaze direction (direct or averted), In addition, we wanted to examine whether there were sex differences when the faces were presented preferentially to the magnocellular versus parvocellular visual pathway. We reported four main findings of sex differences: 1) Female participants recognized the emotion of faces with averted gaze more accurately than did male participants, indicating better ability in females to integrate eye gaze with facial expression, 2) Male participants showed greater right amygdala activation for M-biased averted-gaze fear faces, whereas female participants showed greater involvement of the left amygdala when they viewed both fearful and neutral faces, 3) Female participants had greater amygdala volumes than male participants, with the difference being more pronounced in the right amygdala, and 4) both the left and right amygdala volumes positively correlated with behavioral accuracy for M-biased fear in female participants, whereas only the right amygdala volume correlated with accuracy for M-biased fear in male participants.

When facial expression signals the emotional state of an expresser, eye gaze direction can indicate the source or target of that emotion (Adams & Kleck, 2003; Adams & Kleck, 2005; Adams et al., 2012; Cushing et al., under review; Hadjikhani et al., 2008; Im et al., under review). The social signals conveyed by facial expression and eye gaze can be integrated to facilitate an observer’s processing of emotion of the face when they convey congruent information. For example, processing of approach-oriented facial expressions such as anger or joy can be facilitated with direct eye gaze, which also signals approach motivation, while processing of avoidance-oriented facial expressions, such as fear or sadness, can be facilitated with averted eye gaze which likewise signals avoidance (Adams & Kleck, 2003; 2005; Sander et al., 2007). Therefore, the meaning and intensity of an observed facial expression is driven not only by the expression itself, but also by the changes in an expresser’s eye gaze (Benton, 2010; Cushing et al., under review; Fox et al., 2007; Hadjikhani et al., 2008; Im et al., under review; Milders et al., 2011; N’Diaye, Sander, & Vuilleumier, 2009; Rigato et al., 2013; Sander et al., 2007; Sato et al., 2004). The outcome of the present work further shows that this integration of facial expression and eye gaze is also modulated by the observer’s sex.

Better recognition of emotional faces with averted eye gaze in female than male participants suggests that females are more sensitively tuned to reading and integrating eye gaze with facial expression. Previous studies of sex-related differences in affective processing have also reported female participants outperforming males in facial detection tasks (recognition of a face as a face) or facial identity discrimination and showing stronger face pareidolia, the tendency to perceive non-face stimuli (e.g., food-plate images resembling faces) as faces (Pavlova, Scheffler, & Sokolov, 2015). The superiority of female participants, however, seems to be more pronounced when the task or stimulus involves processing of subtle facial cues, and reduced when highly expressive and obvious stimuli are presented (Hoffmann et al., 2010). Females were also shown to have superior skills in other types of integrative processing during visual social cognition, such as body language reading (e.g., understanding emotions, intentions, motivations, and dispositions of others through their body motion). Specifically, females were faster in discriminating emotional biological motion from neutral and more accurate in recognizing point-light neutral body motion (e.g., walking or jumping on the spot; Alaerts et al., 2011). Our current finding of more accurate integration of facial expression and eye gaze in female observers is in line with these previous findings that showed female observers’ greater ability in integrative and detailed processing of affective stimuli.

In addition to behavioral responses, we also found differences in amygdala activity between female and male observers. Male participants showed greater right amygdala responses, but only to M-biased averted-gaze fear faces (congruent, clear threat cues) whereas the female participants showed greater left amygdala responses to both fearful and neutral faces. Prior work suggests that a fearful face with averted eye gaze tends to be perceived as a clear threat (“pointing with the eyes” to the threat; Hadjikhani et al., 2008) because both emotional expression and eye gaze direction signal congruent avoidance motivation (Adams et al., 2012; Cushing et al., under review, Im et al., under review). Moreover, we previously showed that the right amygdala is highly responsive to such a clear threat cue (averted-gaze fear) (e.g., Adams et al., 2012; Cushing et al., under review), especially when presented to the magnocellular pathway (Im et al., under review). Thus, greater right amygdala reactivity to M-biased averted-gaze fear faces in male participants suggests that processing of facial expression by eye gaze interactions in male observers is tuned more to detection of a clear, congruent threat cue from face stimuli, compared to female observers. Conversely, female observers showed more involvement of the left amygdala, which has been suggested to play a role in more detailed analysis and reflective processing (Adams et al., 2012; Cushing et al., under review; Im et al., under review). Together, our findings suggest that brain activations elicited by interactions of facial expression and eye gaze direction are modulated by the observer’s sex. While sex-specific modulation of fMRI activity for interaction of threatening facial and bodily expressions has been reported (Kret et al., 2011), our study presents the first behavioral and neural evidence of sex-specific modulation of integration of facial expression and eye gaze direction.

Our findings of the differential attunement of female and male observers toward the left and right amygdala are also in line with the previous findings that showed sex-specific hemispheric lateralization of the amygdalae. For example, the two previous studies that involved only male participants showed either exclusive (Cahill et al., 1996) or predominant (Hamann et al., 1999) right lateralization of the amygdala. On the other hand, the two studies that involved only female participants reported left lateralized amygdala activation (Canli et al., 2000, 1999). By directly comparing the amygdala activity between female and male observers in identical conditions, researchers (Cahill et al., 2001) also found a clear difference in hemispheric lateralization of the amygdala in which increased recall of the emotional, compared with neutral, films was significantly predicted by the right amygdala activity in male observers and by the left amygdala activity in female observers. Given the putatively different emphases of the left and right amygdalae in reflective and reflexive processing of compound facial cues (Adams et al., 2012; Cushing et al., under review; Im et al., under review), such sex-related differences in amygdala activity may reflect different cognitive and processing styles in female and male participants, such as better integration of incongruent eye gaze and facial expression cues in females, and a greater bias towards perceiving clear, congruent threat cues in males.

Along with the sex differences in the functional reactivity of the amygdala, we also observed sex-related differences in amygdala volumes. Previous clinical studies have reported that the amygdala volume was correlated with observers’ ability to recognize happy facial expression in Huntington’s Disease (Kipps et al., 2007) and with observers’ anxiety or depression level both in children and in adults (Barrós-Loscertales et al., 2006; De Bellis et al., 2000; Frodl et al., 2002; MacMillan et al., 2003). A recent study of healthy participants has also reported that amygdala volume correlated with individuals’ social network size and complexity (Bickart et al., 2011). Using a large sample of healthy participants, we found that their amygdala volumes also positively correlated with observers’ accuracy for M-biased averted-gaze fear faces (clear threat). This is consistent with previously reported evidence of magnocellular projections to the amygdala for fast, but coarse, threat-related signals (e.g., Méndez-bértolo et al., 2016), and enhanced right amygdala activation for M-biased neutral stimuli (object drawings), compared with P-biased stimuli (Kveraga et al., 2007). Furthermore, we also observed sex-related lateralization differences in that both the left and right amygdala volumes predicted the behavioral accuracy in recognizing M-biased averted-gaze fear faces in female observers, whereas only the right amygdala showed such a relationship in male observers.

## Conclusions

The current study found that an observers’ sex affects behavioral responses to facial expression and eye gaze interaction, and predicts both functional reactivity and volume of the amygdala. Many diseases related to impairments in visual social cognition show sex differences in rates of affliction: Females are more often affected by anxiety disorders (Mclean et al., 2011) and depression (Abate, 2013), whereas males have a higher risk of developing autism spectrum disorders (Werling & Geschwind, 2013) and attention deficit hyperactivity disorders (Ramtekkar et al., 2010). Thus, investigating sex-related differences in behavioral and neural responses, as well as structure, in larger samples of healthy observers will have important implications for identifying such sex differences in these psychiatric disorders. The present findings demonstrate that neural mechanisms underlying affective visual processing can differ between healthy men and women. The current findings further indicate that theories of behavioral and neural mechanisms underlying the perception of affective stimuli should take into account participants’ sex.

## Acknowledgments

This work was supported by the National Institutes of Health R01MH101194 to K.K. and to R.B.A., Jr.

Kestas Kveraga

Reginald B. Adams, Jr.

## Author Contributions

R. B. Adams, and K. Kveraga developed the study concept and designed the study. Testing and data collection were performed by H. Y. Im, N. Ward, C. A. Cushing, and J. Boshyan. H. Y. Im analyzed the data and all the authors contributed to the writing of the manuscript.

